# MacroH2A-Mediated Gene Repression through Nucleosome Compaction and Remodeling Inhibition

**DOI:** 10.64898/2026.01.26.701634

**Authors:** Vladyslava Sokolova, Rulan Jiang, Amber Mullins, Gahyun Lee, Bang Hua Pan, Dongyan Tan

**Affiliations:** Department of Pharmacological Sciences, Stony Brook University; Stony Brook, NY, USA; Program of Pharmacology, Yale University, New Haven, CT, USA; Department of Molecular Biology, Princeton University, NJ, USA

**Keywords:** histone variant, macroH2A, nucleosome, chromatin, chromatin remodeling

## Abstract

MacroH2A (mH2A) is a histone variant primarily implicated in heterochromatin maintenance and transcriptional repression, yet how its conserved histone-fold and its variant-specific domains reshape chromatin to enforce gene silencing remains poorly understood. Here, we dissect the domain-specific contributions of mH2A to nucleosome dynamics and chromatin remodeling. We show that, in addition to its linker region, the C-terminal tail of mH2A histone-fold also stabilizes nucleosome entry/exit DNA. Cryo-EM analysis reveals that this C-terminus tracks along nucleosomal DNA toward the dyad, adopting an on-dyad binding mode reminiscent of linker histone H1. Additionally, the mH2A linker potently inhibits DNA translocation by both the INO80 and Chd1 chromatin remodelers, while the histone fold selectively suppresses INO80 activity. In contrast, the macro domain has no detectable impact on terminal DNA accessibility or remodeling. Together, our results uncover a previously unappreciated architectural role for the mH2A histone fold and establish the linker domain as a dominant regulator of nucleosome dynamics and chromatin remodeling, providing a mechanistic framework for how mH2A enforces transcriptional repression.

## INTRODUCTION

In eukaryotes, the genome is packaged into a DNA–protein complex known as chromatin. The fundamental repeating unit of chromatin is the nucleosome core particle (NCP), which consists of 147 base pairs (bp) of DNA wrapped around an octameric protein core. This core is composed of two copies each of four core histone proteins: H2A, H2B, H3, and H4 (1). The nucleosome is further stabilized by the linker histone H1 that binds at the entry/exit DNA. Histones are among the most conserved proteins in eukaryotes. Their mutation is often lethal, and their dysregulation is associated with various diseases (2).

Nonallelic isoforms of canonical histones, known as histone variants, are incorporated into chromatin in a replication-independent manner throughout the cell cycle. Histone variants confer unique chemical and physical properties to nucleosomes, linking them to specific roles in transcription and genome maintenance (2,3). Among canonical histones, the H2A family exhibit the highest sequence divergence and the largest number of known variants. One such variant, macroH2A (mH2A), is a unique histone best known for its role in the inactivation of X chromosome in female mammals, a specialized domain of heterochromatin (4). Variant mH2A is also frequently found concentrated at other facultative heterochromatin (5,6) and its presence creates a barrier of iPSC reprogramming (7). Moreover, gene bodies of actively transcribed genes are typically depleted of mH2A (8). In sum, existing data support a primarily repressive role for mH2A in transcription (9), with a few exception indicating a gene activation role under certain stress conditions (6,10,11).

Approximately three times the size of its canonical counterpart, mH2A consists of three functional domains: an N-terminal histone-fold domain, a flexible linker region, and a globular C-terminal macro domain (**Fig. 1A**). The disordered linker is highly basic and rich in lysine and arginine residues, resembling the C-terminal tail of linker histone H1. The macro domain can bind NAD+ metabolites (12) and ADP-ribose, with implication in DNA damage repair, but appears to be dispensable for most transcription repression function of the variant. *In vitro*, the mH2A linker has been shown to enhance nucleosome compaction by protecting the DNA near the nucleosome entry/exit sites (13) and promotes nucleosome array condensation (14). The same study showed that the macro domain, in the contrary, inhibits both functions (14). In a separate study, mH2A was shown to interfere with chromatin remodeling by the SWI/SNF complex through its H2A-like histone domain(15).

**Figure 1.**
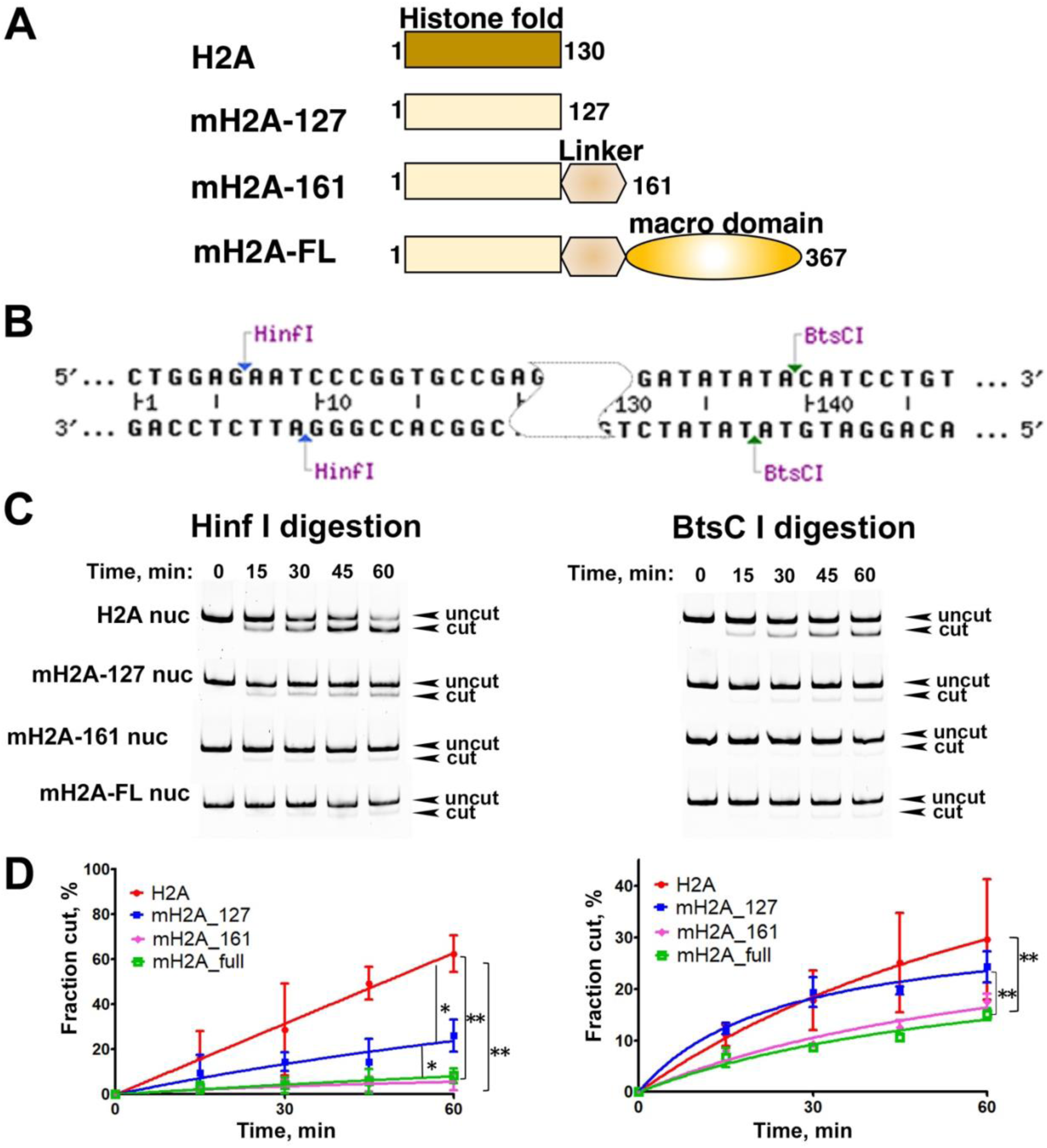
MacroH2A-linker protects Entry/Exit DNA of nucleosomes. A. Representation of the protein domains, used in this study. B. The 601 Widom DNA with the two restriction-enzyme cutting site indicated. C. Representative acrylamide gels of the endonuclease assay of canonical and different truncated mH2A nucleosomes. D. Quantification of the digestion shown in (C). The left panel is the HinfI digestion result and the right is BtsCI digestion results. Data are mean ± SD, n = 3. Statistical significance was indicated by asterisk.

Despite these findings, the precise mechanisms underlying mH2A-mediated transcriptional repression and the distinct contributions of its individual domains remain unclear. To address these questions, we employed cryo-electron microscopy (cryo-EM) and biochemical assays to investigate the effects of mH2A domains on nucleosome structural dynamics and ATP-dependent chromatin remodeling. We present the cryo-EM structure of an mH2A-linker nucleosome stabilized by single-chain variable fragment (scFv) antibodies. The structure reveals a histone fold domain of mH2A closely resembled that of the canonical nucleosome. Although the majority of the flexible mH2A linker remains unresolved in the EM map, a small segment is visible sandwiched between superhelical location 0 (SHL0) and the entry/exit DNA. Our biochemical data show that the mH2A linker interacts with entry/exit DNA but does not engage the distal linker DNA as H1 does. Furthermore, we demonstrate that both the histone domain and the linker of mH2A interfere with ATP-dependent DNA translocation by the INO80 chromatin remodeler. However, only the mH2A linker, not its histone domain, inhibits remodeling by the Chd1 remodeler. The macro domain shows no detectable effect on either DNA protection or nucleosome sliding. Collectively, these results provide mechanistic insights into how individual mH2A domains coordinate to regulate chromatin structure and contribute to transcriptional repression.

## MATERIALS AND METHODS

### DNAs

Three 601 Widom DNAs with different extra-nucleosomal DNA lengths were used in this study, two center-position DNA of 167-bp and 208-bp long and an end-position DNA called 0N80. The DNA template containing twelve tandem repeats of 167-bp 601 Widom sequence was a kind gift from Dr. Craig Peterson. The DNA template containing twelve tandem repeats of 208-bp 601 Widom sequence was a kind gift from Dr. Ed Luk. Large-scale plasmids were purified as previously described (1). A single copy of end-positioned 0N80 (80 base pairs of extra-nucleosomal DNA at one entry/exit site) Widom DNA (WT) was amplified by PCR using a template of plasmid pGEM-3z/601, a kind gift from Jonathan Widom (Addgene plasmid #26656). Multiple copies of the fragment where then cloned into the pUC16 plasmid. To generate a single repeat of these 601 sequences, restriction digestion by EcoRV and ScaI were used to excise the plasmids. DNA fragments were further purified by polyethylene glycol precipitation followed by MonoQ anion exchange chromatography.

The sequences for the 167-bp, 208-bp, and 0N80 Widom sequence are listed below with the 601 sequence underlined:

167-bp DNA repeat: ATCCCGCCCTGGAGAATCCCGGTGCCGAGGCCGCTCAATTGGTCGTAGACAGCTCTAGCACCGCTTAAACGCACGTACGCGCTGTCCCCCGCGTTTTAACCGCCAAGGGGATTACTCCCTAGTCTCCAGGCACGTGTCAGATATATACATCCTGTGCATGACTAGAT

208-bp DNA repeat: ACTTATGTGATGGACCCTATACGCGGCCGCCCTGGAGAATCCCGGTGCCGAGGCCGCTCAATTGGTCGTAGACAGCTCTAGCACCGCTTAAACGCACGTACGCGCTGTCCCCCGCGTTTTAACCGCCAAGGGGATTACTCCCTAGTCTCCAGGCACGTGTCAGATATATACATCCTGTGCATGTATTGAACAGCGACCTTGCCGGAGT

0N80 DNA repeat: ACTGAGAATCCCGGTGCCGAGGCCGCTCAATTGGTCGTAGACAGCTCTAGCACCGCTTAAACGCACGTACGCGCTGTCCCCCGCGTTTTAACCGCCAAGGGGATTACTCCCTAGTCTCCAGGCACGTGTCAGATATATACATCCTGTGCATGTATTGAACAGCGACCTTGCCGGTGCCAGTCGGATAGTGTTCCGAGCTCCCACTCTAGAGGATCCCCGGGTA CAGT

### Protein production and Nucleosome Reconstitution

The human macroH2A1.1 variant without any affinity tag (mH2A-FL) was used in this study. The mH2A_161_ construct contains the histone fold and the linker region of the protein (residue 1 to 161). The mH2A_127_ construct contains only the histone fold (residue 1 to 127). The mH2A_161_-swap and mH2A_127_-swap mutants were obtained by substituting residues 109-127 in mH2A with residues 112-130 in human H2A.1.

*Xenopus laevis* histones H2A, H2B, and H4 were expressed in BL21 (DE3) *E. coli* cells and H3 was expressed in BL21 (DE3) pLysE *E. coli* cells. The different versions of mH2A variant were expressed in BL21 (DE3) *E. coli* cells. All histones were purified as previously described (16), with the mH2A histones purified using a modified protocol described below. Specifically, the pellet was resuspended in urea unfolding buffer [7 M Urea, 20 mM Tris (pH 7.5), 1 mM EDTA, 500 mM NaCl, and 1 mM DTT] supplemented with protease inhibitors. The lysate was then sonicated, and incubated with additional urea unfolding buffer in a rotator at 4°C for 1 h. The cell lysate was cleared by centrifugation at 15,000×g at 4°C for 45 min and filtered through 1 μm and 0.45 μm filters. The clear lysate was then loaded onto a HiTrap Q HP column (Cytiva) pre-equilibrated with buffer [7 M Urea, 20 mM Tris (pH 8.0), 1 mM EDTA, 500 mM NaCl, and 1 mM DTT]. The flowthrough was collected and diluted with dilution buffer [7 M Urea, 20 mM Tris (pH 8.0), 1 mM EDTA, and 1 mM DTT] before loading onto a HiTrap S HP column (Cytiva). The column was first washed with buffer A [7 M Urea, 20 mM Tris (pH 8.0), 1 mM EDTA, 100 mM NaCl, and 1 mM DTT], followed by protein elusion using a gradual gradient between buffer A and buffer B [6.2 M Urea, 20 mM Tris (pH 8.0), 1 mM EDTA, 1 M NaCl, and 1 mM DTT]. Eluted fractions were analyzed by SDS-PAGE and the best fractions were collected and dialyzed into water.

To reconstitute histone dimer and tetramer, an equimolar of the corresponding histones were mixed. The mixture was incubated for 2 h in unfolding buffer [7 M guanidine HCl, 20 mM Tris (pH 7.5), and 10 mM DTT] followed by dialysis against at least three changes of refolding buffer [10 mM Tris (pH 7.5), 1 mM EDTA, 2 M NaCl and 1 mM DTT] at 4°C. Dimer or tetramer was concentrated and purified via size-exclusion chromatography using a Superdex200 increase 10/300 GL column (Cyntiva).

### Expression and purification of recombinant INO80-C complexes

The INO80 complex used in this study contains 10 subunits of the *S. cerevisiae* complex. It is equivalent to the INO80-C ΔN sub-complex in our previous study (17). The Ino80 motor in this sub-complex is a truncated form of the Ino80 subunit (residue 470 from 1490). The complex was generated using the MultiBac system (Geneva Biotech) and expressed and purified as described previously (17).

### Nucleosome Reconstitution

To reconstitute nucleosomes, histone dimer, tetramer, and DNA (167-bp, 208-bp, or 0N80 DNA) were mixed in high-salt buffer [10 mM Tris (pH 8.0), 2 mM EDTA, 2 M NaCl, and 2 mM 2-mercaptoethanol (βME)]. The mixture was dialyzed overnight into low-salt buffer [10 mM Tris (pH 8.0), 2 mM EDTA, 5 mM NaCl, and 2 mM βME]. The optimal ratio of dimer to tetramer to DNA was determined empirically through titration and examination via electromobility shift assays (EMSA).

### scFv antibody production

A single-chain fragment variable (scFv) antibody plasmid was a kind gift from Dr. Yawen Bai and was expressed and purified essentially as described with minor modifications (18). Briefly, scFv was expressed in Bl21(DE) cells at 25℃ for 16h. Cells were harvested and resuspended in Lysis buffer (50 mM Tris-HCL, pH7.5, 100 mM NaCl, 1 mM EDTA). Cells were disrupted by sonication. Inclusion body was pelleted by centrifugations (15000xg for 45 min), washed with lysis buffer and pelleted again. Inclusion body was resuspended in denaturing buffer (100 mM Tris-HCl pH8, 6 M Guanidine, 2mM EDTA, 10 mg/ml DTE), followed by sonication and rotation in the cold room overnight and a final centrifugation (15000xg for 45 m). Antibody was refolded by quickly adding 10 ml of the clear extract from centrifugation to 1 L of refolding buffer (100 mM Tris-HCl, 1mM EDTA 0.5 M arginine, pH9.5, with added freshly oxidized glutathione 551 mg/L) with 3 min stirring. Refolding solution was incubated at 10℃ for 48h without stirring, followed by dialysis at 4℃ against two changes of dialysis buffer (20 mM Tris–HCl, pH 7.4, 100 mM urea freshly added) for 16 h each. The solution was cleared by centrifugation and filtration, and then subjected to further purification using HiTrap SP HP cation exchange column (Cytiva). Fractions containing scFV were pooled and subjected to affinity purification using His-trap column (Cytiva), followed by size-exclusion chromatography using Superdex200 increase 10/300 GL column. Pure fractions were pooled, concentrated (final 5uM), and stored on ice until use.

### Endonuclease digestion Assay

250 nM of the 167-bp nucleosomes were mixed with either 45 U of the HinfI endonuclease or 4.5 U of the BtsCI endonuclease in Cutsmart buffer (NEB) [20 mM Tris-Ac (pH 7.9), 50 mM KAc, 10 mM MgAc, and 100 µg/mL BSA]. The total volume of each reaction was 25 µL. The reactions were incubated at 37 °C and 5 µL sample was collected every 15 min. The reaction was quenched by adding 8 µL stop buffer [10 mM Tris–HCl (pH 8.0), 0.6% SDS, 40 mM EDTA, and 0.1 mg/mL proteinase K], followed by deproteination at 50°C for 1 h. The samples were then separation on 8% Native-PAGE gels (100 V, 180 min, 1x TBE). The gels were stained with SYBR-GOLD (GoldBio) and imaged on a Typhoon imager (Cytiva). The gels were quantified using the ImageJ software version 1.53e. The level of significant difference was determined using a two-way ANOVA test, with *p* ≤ 0.05 being considered significant. The graphical representations and two-way ANOVA tests were performed using Prism 5 software.

### MNase digestion Assay

425 nM of 167-bp nucleosomes (with scFv-bound) were subjected to digestion with 0.75U of MNase (Roche) in MNase buffer (10mM Tris pH7.4, 50mM NaCl, 2mM CaCl_2_, 0.1mg/ml BSA) at 37℃. 4.5 µL of sample was collected every 3 min with the reaction quenched by adding 10 µL of stop buffer [10 mM Tris (pH 7.5), 40 mM EDTA, 0.6% SDS, 0.1mg/ml proteinase K] and incubated at 50°C for 1 h. Samples were separated on an 8% Native-PAGE with 2.5% stacking gel (19:1Acrylamide/Bis) at 4°C (100 V, 180 min, 1x TBE buffer). The gels were stained with SYBR-GOLD for 10 min. Gels were imaged on a Typhoon imager (Cytiva). Quantification of the gels was done using ImageJ software. Intensities for DNA fragments were estimated cumulatively from bands of similar size. The two-way ANOVA test was used to determine whether the differences between data sets are statistically significant using the p ≤ 0.05 criterion. Two-way ANOVA test and graphical representation were done using Prism 5 software.

### Nucleosome-sliding assay

End positioned 0N80 nucleosomes (140 nM) were incubated for 10 min with recombinant yeast INO80 complex or commercial Chd1 protein (Active Motif, Catalog No: 81307) (50 nM) in sliding buffer [25 mM HEPES (pH 8.0), 50 mM NaCl, 2.5% glycerol, 1 mM TCEP, and 2 mM MgCl_2_]. Reaction was initiated by adding 1 mM ATP and quenched by adding stop-buffer [5 mM EDTA, 0.2 mg/mL lambda DNA (NEB), and glycerol 10%]. Sliding by INO80 was performed at room temperature, and sliding by Chd1 was performed at 30°C. After quenching for 10 minutes all samples were resolved on 6% Native-PAGE at 4°C for 2 h (for the INO80 experiment) or 3 h (for the Chd1 experiment). Gels were stained with SYBR-GOLD before imaging on a Typhoon imager (GE Healthcare).

### Assembly of the scFv-mH2A_161_ nucleosome complex for cryo-EM study

mH2A_161_ nucleosomes (containing 208-bp of 601 Widom DNA sequence) were combined with approximately 2-fold of scFv in low-salt buffer [10 mM Tris (pH 8.0), 2 mM EDTA, 5 mM NaCl, and 2 mM DTT]. The sample was concentrated and further purified using Superose 6 Increase 10/300 Gl column in TEN10 buffer [10 mM Tris (pH 7.5), 1 mM EDTA, 10 mM NaCl and 0.5 mM TCEP] as described (18). Fractions were analyzed by 4% Native-PAGE at 4°C (100 V, 75 min, 1× TBE). Good fractions were pooled and concentrated to ∼5 µM. The sample was cross-linked with 0.1% glutaraldehyde for 15 minutes on ice and the reaction was stopped by adding Tris (pH 7.5) to a final concentration of 80 mM.

### Cryo-EM sample preparation and imaging

For the macroH2A_161_-scFv nucleosome, grids were vitrified using Vitrobot Mark IV (FEI Company) operated at 4℃ and 95% humidity. 3.5 µl specimen were applied to glow-discharged QUANTIFOIL grids (R1.2/1.3 – 400 mesh). The grids were then blotted for 4.5-5 seconds with 0 blot force, before plunged frozen in liquid ethane. The frozen grids were stored in liquid nitrogen until screening and data collection. Screening was done in the cryo-EM facility in Stony Brook University using the 200kV Talos Arctica microscope (FEI). Data collection was performed at the Laboratory of BioMolecular Structure (LBMS) in the Brookhaven National Laboratory. Data were acquired on a FEI Titan Krios microscope operated at 300 kV with a nominal magnification of 105,000X and a Bioquantum energy filter (Gatan) operating at zero loss frequency 10 eV. Movies were recorded using a K3 direct detector (Gatan) in super-resolution mode using EPU software, giving a pixel size of 0.4125 Å at the specimen level. Defocus ranges from - 1.0 to -2.25 µm. Each movie was dose-fractionated to 40 frames with a dose rate of ∼1.25 e/Å^2^/sec. Total dose per micrograph is 50 e/Å^2^.

### Image analysis

Multi-frame movies were aligned and summed using the motion correction software implemented within RELION (19). The CTF parameters were estimated using CTFFIND4 (20). Particle-picking was carried out using Topaz neural-network picking (21). Bad particles were removed through multiple rounds of 2D classifications, resulting in a data set of 1.9-millions particles. These data were then subjected to 3D classification in RELION. To generate an initial model, selected 2D classes representing various views of the complex were chosen and used for the Ab-Initio reconstruction in RELION. The subsequent low-resolution map was used as the initial model for 3D classification in RELION. After three rounds of 3D classifications, the best class containing 448,531 particles was selected and subjected to consensus 3D refinement, generating a density map with average resolution of 3.0 Å. CTF refinement and postprocessing was performed, which improved the resolution to 2.8 Å. The global resolution of the map was estimated using the gold-standard Fourier Shell Correlation (FSC) 0.143 criterion with automatic B factor determined in RELION. Local resolution was estimated in RELION.

To resolve the disordered mH2A C-terminal tail and the linker, we performed a DynaMight job in RELION and obtained a deformed 3D reconstruction with improved densities at the mH2A C-terminal tail and the entry/exit DNA. Figures were generated using UCSF ChimeraX (22) and Coot (23). Movies were created using UCSF ChimeraX.

### Alphafold3 Prediction

The AlphaFold3 predicted structure of human mH2A1.1 full-length nucleosome and mH2A_161_ nucleosome were visualized using ChimeraX (22). Both nucleosomes also contain canonical Xenopus histone H2B, H3, and H4.

### Model Building

The AlphaFold3 predicted structure of mH2A_161_ nucleosome, the crystal structure of scFv (PDB: 7k5x), and 601 DNA (PDB: 3lz0) were rigid-body fitted to the cryo-EM density map using UCSF ChimeraX to generate an initial model. All flexible histone tails and the mH2A linker were removed in the initial model as their corresponding densities were not resolved in the map. This model was then subjected to global real-space refinement and model minimization in PHENIX (24). The final structure was assessed by Molprobity (25). Model visualization and figures generation were carried out using the PyMOL Molecular Graphics System (Schrödinger, LLC).

## RESULTS

### mH2A linker is the main contributor for entry/exit DNAs protection on nucleosomes

To determine the contribution of mH2A domains in the mH2A-mediated regulation of nucleosome dynamics and chromatin functions, we generated three constructs of the human mH2A1.1 variant: full-length mH2A (mH2A-FL), mH2A containing the histone fold and its linker region (mH2A_161_), and mH2A histone fold only (mH2A_127_) (**Fig. 1A**). These variation of mH2A variant alongside *Xenopus* histones H2B, H3, and H4 and the 601 Widom DNA were used to reconstitute mono-nucleosomes (**Fig. S1B-C**).

Nucleosomes inherently exhibit entry/exit DNA breathing as an intrinsic property. Certain histone variants, such as H2A.Z and H2A.B, enhance nucleosome DNA breathing and increase their accessibility (26,27). Variant mH2A has the opposite effect, shown by an early study where its linker prevents entry/exit DNA from exonuclease III digestion (13). In our study, both mH2A-FL and mH2A_161_ nucleosomes inhibit nucleosomes digestion by restriction enzymes HinfI and BstCI to a similar degree (**Fig 1B-D**). The mH2A_127_ nucleosome, on the other hand, behaved more similarly to the canonical nucleosome. However, it exhibited differential cleavage, where no difference was observed between mH2A_127_ and canonical nucleosomes upon BstCI treatment (**Fig. 1C&D**), while mH2A_127_ inhibited HinfI digestion with a modest but significant level, albeit the effect was less pronounced than with mH2A-FL and mH2A161 nucleosomes. These results support previous findings that mH2A linker interacts and protects terminal DNA on nucleosomes while the macro domain does not contribute to additional DNA protection nor counteract such effect.

We next asked if DNA methylation, a modification generally associated with gene repression, synergizes with mH2A-mediated nucleosome DNA compaction. We found that HinfI digestion of H2A nucleosomes assembled on CpG-methylated 601 DNA was significantly faster than digestion of canonical H2A nucleosomes assembled on unmethylated DNA. A similar trend was observed for mH2A161-containing nucleosomes, although the difference between methylated and unmethylated substrates was more modest (**Fig. S2**). These observations are consistent with recent reports demonstrating that DNA methylation alters nucleosome stability in a histone variant–dependent manner, including effects on H2A.Z-containing nucleosomes. (28).

### The effects of mH2A-linker and H1 in stabilizing linker DNAs

To test whether mH2A linker acts in a similar manner as the globular domain of H1 to compact and protect linker DNAs, we performed micrococcal nuclease (MNase) digestion assays to delineate the DNA binding footprints of the mH2A linker (**Fig 2A**). This assay enables us to evaluate end-DNA protection in both mH2A_161_ and canonical H2A nucleosome, with and without linker histone H1. Specifically, we compared MNase profiles of four reconstituted nucleosomes: H2A nucleosome, H2A nucleosome with H1 (chromatosome), mH2A_161_ nucleosome, and mH2A_161_ nucleosome with H1 (mH2A chromatosome).

**Figure 2.**
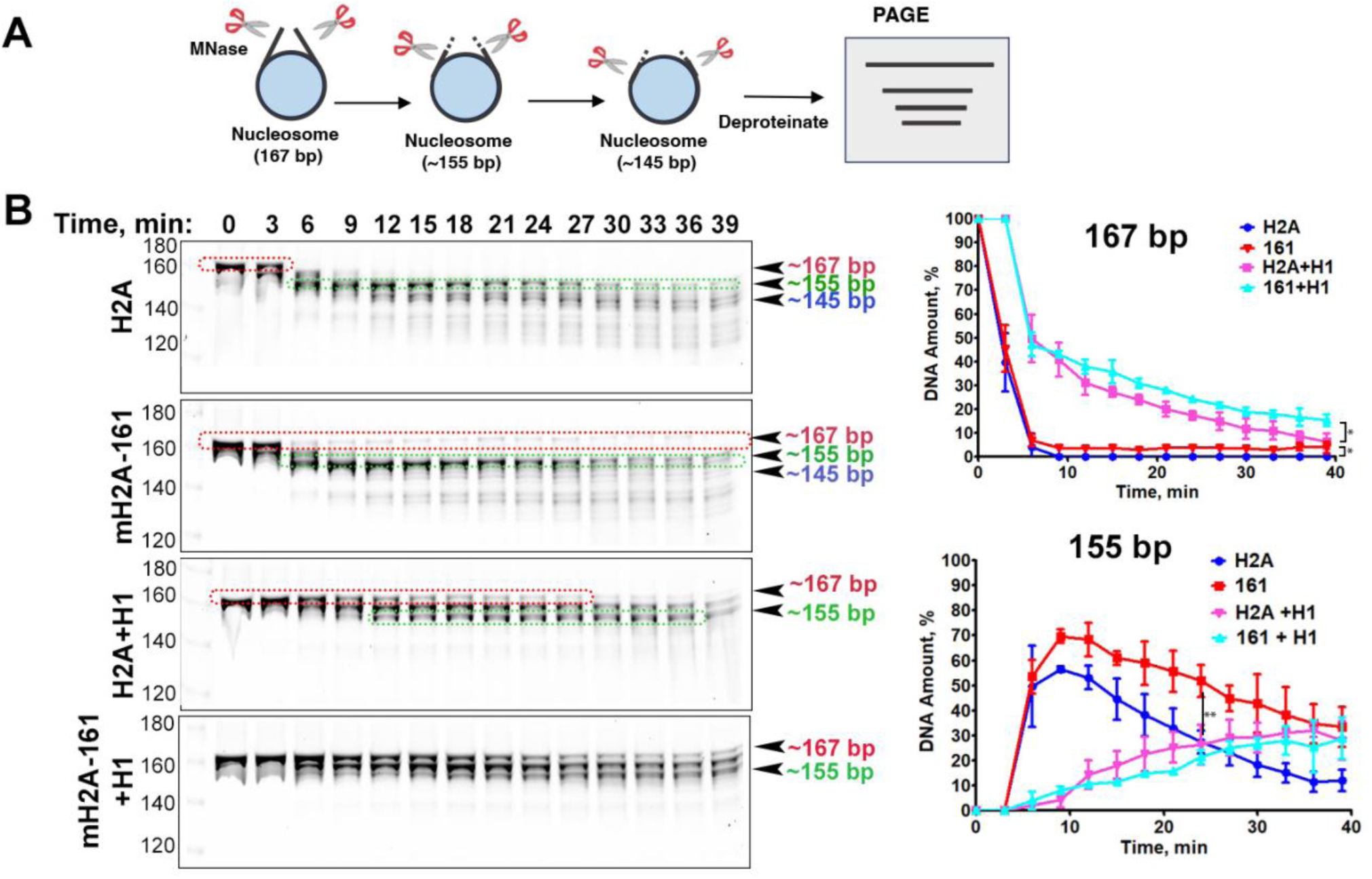
The effect of mH2A_161_ on DNA protection in comparison with linker histone H1. A. Representative acrylamide gels of the MNase assay of canonical, macroH2A-161 nucleosomes with and without addition of linker histon H1. The digestion products of different sizes are labeled. B. Quantification of the MNase assay based on 167-bp and 155-bp bands. Each graph contains all four different nucleosomes. The graph shows the fraction of digested nucleosomes as a function of time. The data points are the average of different biological replicas. Data are mean ± SD, n = 3. Statistical significance are labeled with an asterisk.

As abovementioned, the globular domain of H1 isoforms binds at the nucleosome dyad, contacting both the nucleosomal DNA and the proximal linker DNAs, while its C-terminal tail interacts with the distal linker DNAs to modulate their conformation (18). Consistent with these known interactions, MNase digestion of human H1.0-containing chromatosomes proceeded much more slowly than in either mH2A_161_ or H2A nucleosomes. Quantification revealed that the 167-bp DNA fragment was completely digested within 5 minutes in both mH2A_161_ and H2A nucleosomes, but remained detectable in chromatosomes even after 30 minutes (**Fig. 2B**). This result is consistent with previous findings showing that full-length H1 shields additional 20-bp of linker DNA. In addition, two shorter DNA fragments with size of 155 bp and 145 bp, were observed. In both chromatosomes, the increase of 155-bp fragment correlates with the decrease of 167-bp fragment during the course of the experiments (**Fig 2B**), in line with the protection of H1 on linker DNAs.

We also detected significant differences between the digestion patterns of mH2A_161_ and canonical H2A nucleosomes. Specifically, mH2A_161_ nucleosomes were digested more slowly than H2A nucleosomes, with a greater retention of the 155-bp fragment. These results indicate that the mH2A linker confers modest but significant protection of entry/exit DNA (**Fig 2B**), but not the distal part of the linker DNA like H1. The fact that mH2A_161_ nucleosome containing H1 show nearly identical level of linker DNA protection compared to chromatosome in our assay suggest a non-overlapping binding mode of the mH2A_161_ linker and H1 on DNA.

### MacroH2A inhibits DNA translocation by INO80 and Chd1 in distinct manners

Nucleosomes containing the histone variant mH2A have been linked to inhibition of chromatin remodeling, but the limited studies available have reported conflicting findings. An early study reported that the non-histone region of mH2A hinders transcription factor binding, while the histone domain interferes with nucleosome remodeling by the SWI/SNF complex (15). In contrast, subsequent work suggested that the non-histone region alone is responsible for inhibiting transcription initiation and chromatin remodeling by SWI/SNF and ACF (29). Adding further complexity, another biochemical study reported that mH2A-containing nucleosomes are actually efficient substrates for both SWI/SNF and ISWI remodeling enzymes (30). These discrepancies underscore the need to clarify which domains of mH2A mediate remodeling inhibition and how they function cooperatively.

To address these questions and dissect the domain-specific contribution of mH2A to chromatin remodeling, we performed nucleosome sliding assays based on electrophoretic mobility shift analysis (EMSA). We examined the effects of truncated mH2A variant on ATP-dependent nucleosome remodeling by two remodelers: the INO80 complex and Chd1. INO80 is a ∼1 MDa, multi-subunit complex composed of 15 subunits, while Chd1 is a single-subunit remodeler. Both catalyze ATP-dependent DNA translocation but engage nucleosomes differently: Chd1 binds at superhelical location 2 (SHL2) on nucleosomal DNA (31), whereas INO80 adopts a dual-binding mode at SHL-2 and SHL-6 positions (32,33).

Our results showed that nucleosomes reconstituted with either mH2A-FL and mH2A_161_ strongly inhibit nucleosome sliding by INO80 (**Fig. 3A &B**) and Chd1 (**Fig. 3C &D**). Intriguingly, mH2A_127_ nucleosomes also inhibited INO80-mediated nucleosome sliding, though the effect was significantly weaker compared to mH2A_161_ and mH2A-FL nucleosomes (**Fig. 3A &B)**. In contrast, mH2A_127_ nucleosome did not impair Chd1-mediated remodeling (**Fig. 3C &D**). These findings indicate that the mH2A linker is the primary determinant of remodeling inhibition, while mH2A histone-fold domain contributes modestly and specifically to INO80 inhibition. The macro domain does not appear to contribute to remodeling inhibition nor to counteract the mH2A linker’s inhibitory effect.

**Figure 3.**
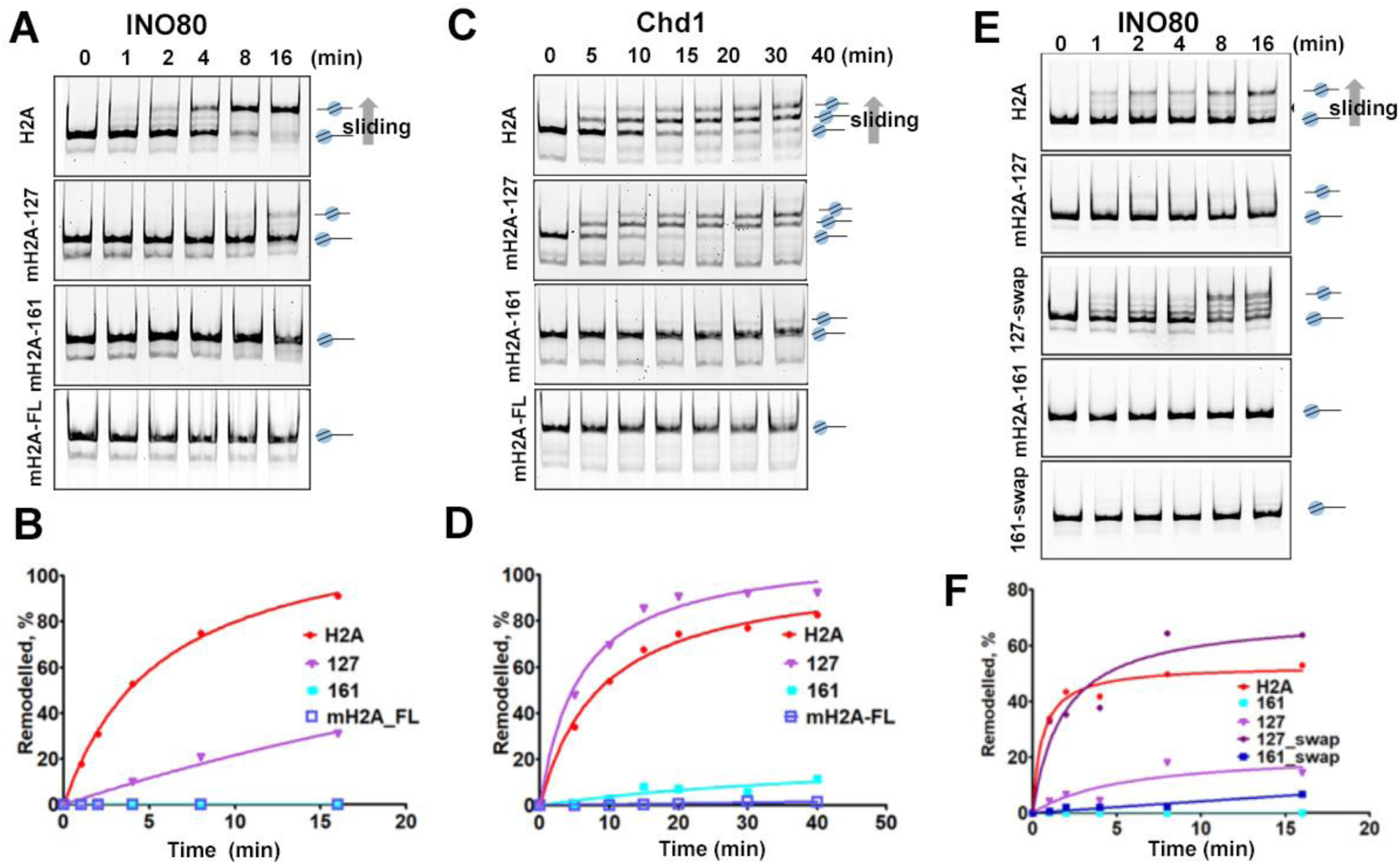
Effects of mH2A variant on INO80 and Chd1-mediated nucleosome sliding. (A) Native-PAGE shows nucleosome-sliding mediated by INO80_core_ complex on H2A, mH2A_127_ and mH2A_161_ nucleosomes, respectively. The bands for end-positioned and center-positioned nucleosomes are indicated by schematics on the right side of the gels. (B) Quantification of the experiment in (A). (C) Native-PAGE shows nucleosome-sliding mediated by Chd1 on H2A, mH2A_127_ and mH2A_161_ nucleosomes, respectively. End-positioned and remodeled nucleosomes are indicated by schematics on the right side of the gels. (D) Quantification of the experiment in (C). (E) Native-PAGE showing nucleosome-sliding in wild-type and two different mH2A chimera mutant nucleosomes. (F) Quantification of the experiment in (E).

### The C-terminus of mH2A histone-fold contribute to remodeling inhibition

The C-terminus of histone fold domain in the H2A family protein, encompassing approximately the majority of the docking domain and the last 20 residues, is among the most sequence-divergent regions. We previously showed that this region in H2A.Z promoted nucleosome destabilization through enhanced terminal DNA flexibility (26). In contrast to H2A.Z, mH2A retains the canonical acidic patch but differs from H2A primarily in the last ∼20 residues of its histone-fold domain (referred as C-terminal tail in **Fig S1A**). To test if these 20-residue segment contributes to mH2A-mediated terminal DNA stabilization and INO80 remodeling inhibition, we generated two chimeric mH2A proteins, mH2A_127-swap_ and mH2A_161-swap,_ in which the last 19 residues of mH2A histone fold were replaced by their counterparts in H2A. Restriction enzyme digestion assays showed that both chimeric mH2A nucleosomes exhibited digestion patterns similar to their wild-type counterparts when treated with HinfI and BtsCI (**Fig. S3A&B**), suggesting that the C-terminal tail does not significantly affect entry/exit DNA accessibility.

We next examined whether the chimeric mH2A nucleosomes differed in their response to INO80-mediated remodeling. While H2A.Z nucleosomes are known to be more efficiently remodeled by INO80 than canonical H2A (34), the responsible feature remains unclear. As shown earlier, the disordered mH2A linker restricts DNA accessibility and inhibits both INO80 and Chd1 activities. To investigate the possible effect of the C-terminal tail, we performed INO80 sliding assays using nucleosomes containing the mH2A swap-mutants. Notably, mH2A_127-swap_ nucleosomes were efficiently remodeled by INO80, resembling H2A nucleosomes (**Fig 3E&F**). In contrast, mH2A_161-swap_ nucleosomes showed no improvement over wild-type mH2A_161_ nucleosomes, likely due to the persistent presence of the inhibitory mH2A linker. Since INO80 binding assays showed comparable affinity for both chimeric and wild-type mH2A nucleosomes (**Fig S4**), we conclude that the reduced INO80 remodeling activity on mH2A nucleosomes is not due to impaired binding. Together, these finding indicates that while the C-terminal tail in the mH2A histone-fold alone do not significantly contribute to mH2A-mediated terminal DNA protection, they inhibit INO80-mediated nucleosome sliding.

### Cryo-EM structure of scFv-mH2A_161_ nucleosome

To gain insights into mH2A linker-mediated linker DNA protection, we performed single-particle cryo-EM study on the mH2A_161_ nucleosome (**Fig S5, Suppl Table 1 &2**). A previous study showed that a scFv from the pl2-6 antibody, originally isolated from autoimmune mice, binds to the acidic patch of the nucleosome core particle (NCP), stabilizing nucleosomes and chromaotosomes for structural studies (35). An additional advantage of using scFv to assist cryo-EM study is that it improves orientation distribution of particles in vitreous ice, mitigating the preferred orientation problem commonly encountered in single-particle cryo-EM studies.

Selected 2D class averages revealed multiple views of the scFv-mH2A_161_ nucleosome, with visible secondary structural features. Consensus refinement produced a density map with an average resolution of 2.8Å (**Fig 4A and supplemental movie 1**), showing well-defined side chain densities of the histone cores (**Fig S6A**) and clear base-pairs features of the DNA (**Fig S6B**). The corresponding atomic model revealed a protein core nearly identical to that of the major-type nucleosome (**Fig S7A**). The model shows 149 bp of DNA compared to the 147 bp alpha-satellite 5S DNA in the crystallographic model of mH2A nucleosome (**Fig S7**). The disordered mH2A linker could not be resolved in the density map.

**Figure 4.**
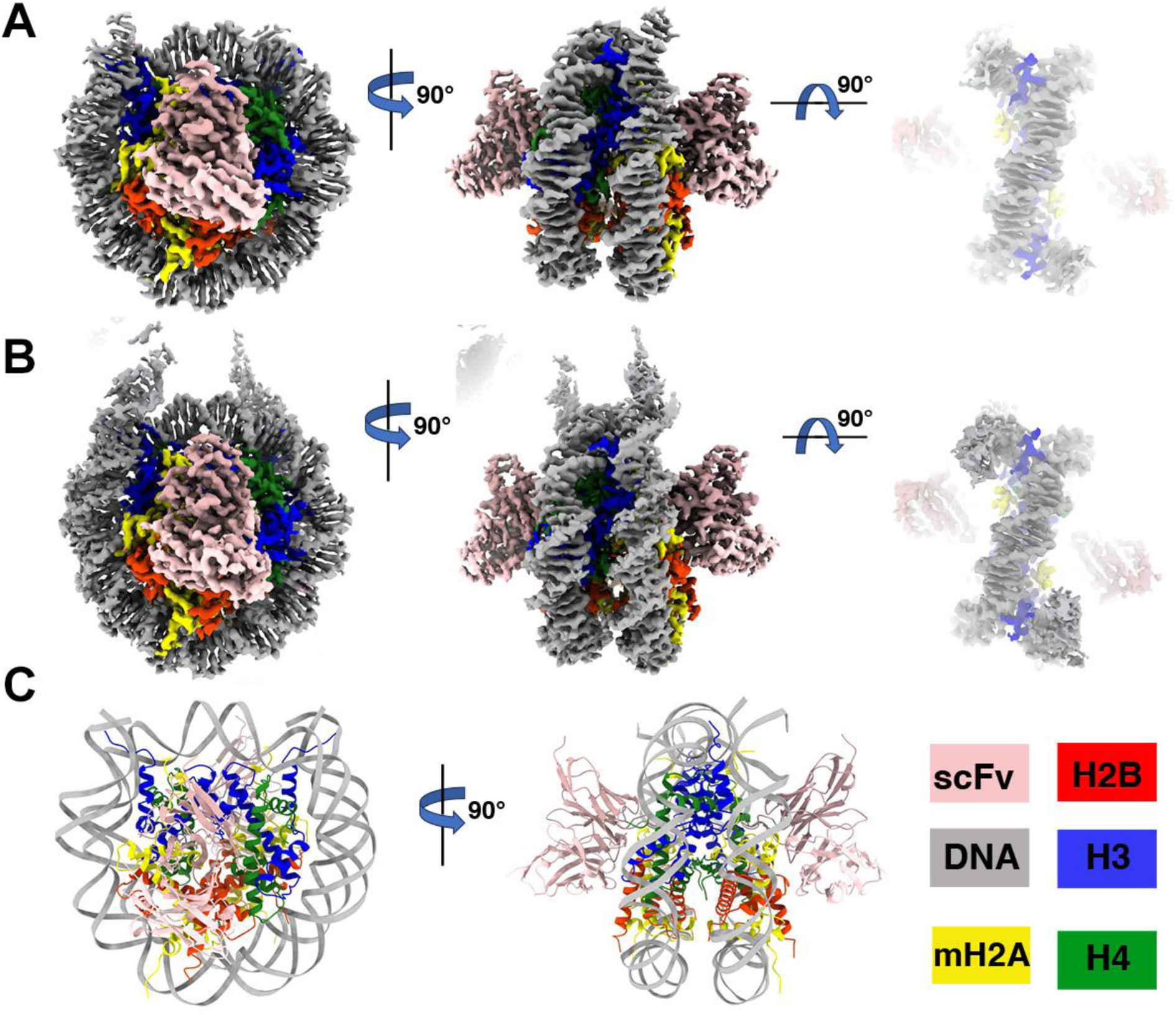
Cryo-EM map and molecular coordinate of mH2A-linker nucleosomes with scFv. A. Postprocess map in three different views, zone color B. DynaMight deformed reconstruction in three different views C. Model of mH2A-161 nucleosome with scFv

In an attempt to resolve the conformational dynamics of mH2A linker, we performed both DynaMight in Relion (36) and 3D Variability analysis in cryoSPARC (37) with the same dataset that produced the consensus refined map. Deformed reconstruction from DynaMight is comparable to the consensus-refined map in terms of resolution, while longer DNA ends are observed in the DynaMight map (**Fig 5B & supplemental movie 2**). Volume series generated from 3D variability analysis also showed the mobility of the complex largely resides in the entry/exit DNA (**supplemental movie 3**).

**Figure 5.**
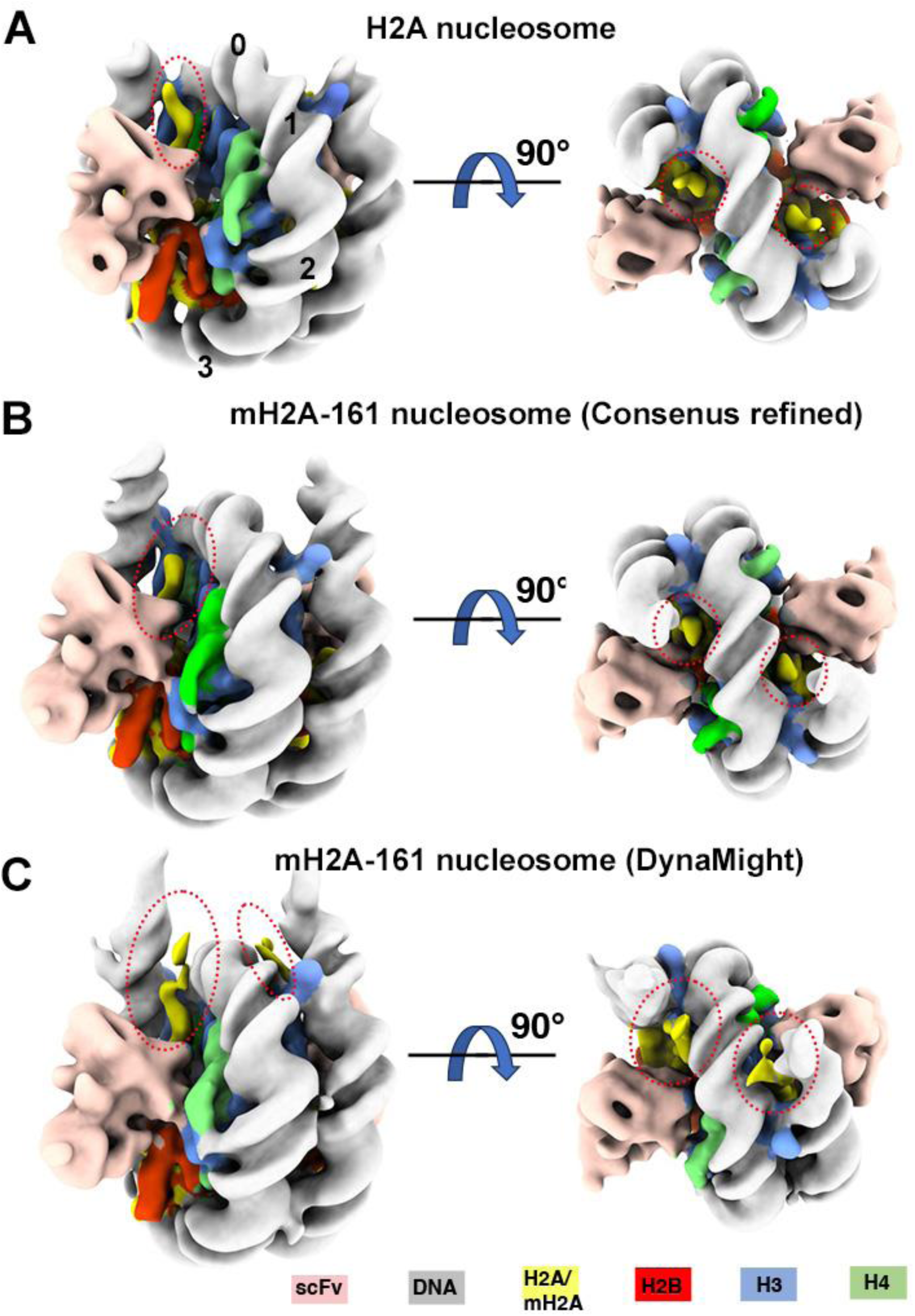
Consensus refinement and dynamite deformed reconstruction of the mH2A-linker nucleosome. A. Canonical nucleosome map low pass filtered to 8 Å B. Consensus refined map low-pass filtered to 8Å. C. DynaMight deformed corrected map low-pass filtered to 8Å

We low-pass filtered both the consensus-refined map and the DynaMight map to 8 Å, and compared them to the published H2A nucleosome-scFv complex to assess whether additional density attributable to mH2A_161_ was present. When all three maps were viewed at comparable contour levels, the DynaMight map revealed additional density representing longer mH2A C-terminal tails that were not typically resolved in cryo-EM maps of nucleosomes (**Fig 5C**). The two C-terminal tails of mH2A in the map are positioned between the entry/exit DNA and SHL0, bending towards the dyad (**Fig 5C**), likely making interactions with the DNA. These observations demonstrated that DynaMight effectively captures the flexible C-terminal tails of mH2A. Although the mH2A_161_ linker region remains disordered, the observed position of the C-terminal tails of mH2A histone fold, combined with prior evidence that the mH2A linker stabilizes the linker DNA (13), led us to hypothesize that the mH2A_161_ linker interacts with DNA in a manner similar to the on-dyad binding mode of linker histone H1 (18). Notably, this configuration contrasts sharply with the Alphafold3 predicted structures of mH2A_161_ and mH2A full-length (FL) nucleosomes, where the linkers project away with no apparent engagement with either DNA or the histone core (**Fig S7B&C**).

## DISCUSSION

Histone variants alters the biochemical and physical properties of chromatin by modifying the nucleosome surface, thereby influencing chromatin compaction and its interactions with epigenetic regulators. In recent years, molecular-level studies have revealed how variant-specific structural features underlie their distinct biological functions. Members of the H2A family are among the most diverged histones, and they also typically exert larger effects on chromatin structures and DNA dynamics (27). For example, both the essential variant H2A.Z and the testis-specific H2A.B possess a truncated C-terminal tails, and such characteristic but not specific amino acids in the sequence have been shown to destabilize DNA wrapping at the nucleosome entry/exit site (38–40).

In this study, we focused on mH2A, the largest and structurally unique histone variants, to gain molecular insights of its role in transcription repression. Our cryo-EM structure reveals that the C-terminal tails of mH2A histone fold domain extend along the entry/exit DNA towards the dyad axis. This result prompt us to propose that the mH2A linker, which follows the C-tail, adopts a similar trajectory to interact and stabilize the linker DNAs (**Fig 6**). We further propose that such arrangement stabilizes the proximal linker DNA, reminiscent of the globular domain of H1 that binds on dyad on nucleosomes (18).

**Figure 6.**
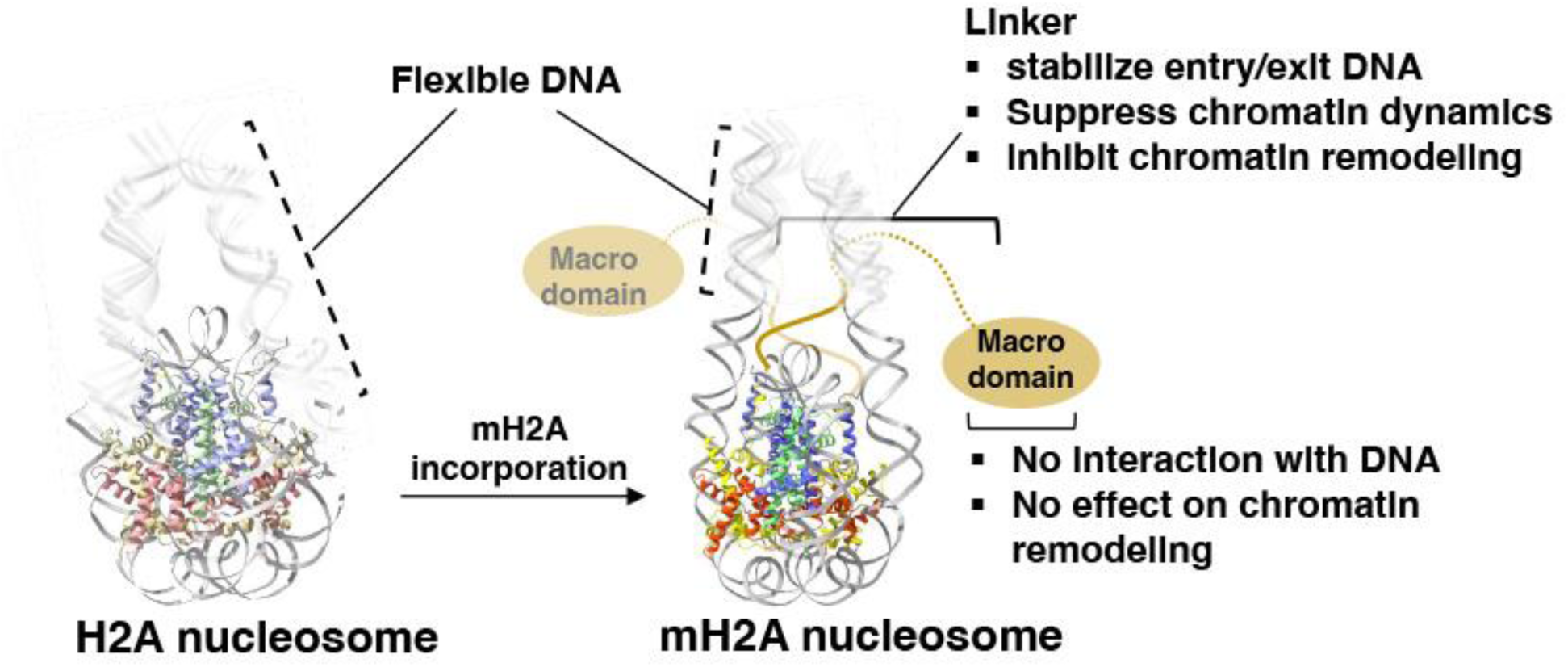
Model of transcription repression by mH2A variant. Incorporation of variant mH2A stabilizes the nucleosome by restricting the entry/exit DNAs through interactions with its disordered linker region. Both the mH2A linker and the C-terminus of its histone-fold can interfere with ATP-dependent chromatin remodeling.

Both our restriction enzyme and MNase digestion assays confirm previous findings, showing that the disordered, lysine-rich mH2A linker plays a dominant role in protecting linker DNA. Unlike earlier reports, our restriction enzyme digestion assays showed that nucleosomes containing only the mH2A histone fold exhibit slight but significant protection of entry/exit DNA compared to H2A nucleosome, at least on one DNA end. For H2A.Z variant, the C-terminal extension of H2A.Z plays a key role in promoting terminal DNA unwrapping and nucleosome destabilization (26). Given that our mutagenesis study revealed no effect of swapping the mH2A C-terminal tail with its canonical counterpar, we speculate that a longer stretch of the mH2A C-terminus, such as its docking domain, are needed to mediate end-DNA protecting through the mH2A histone-fold.

Our study showed that the mH2A C-terminal tail is capable of impeding INO80-mediated nucleosome sliding, as revealed by mutagenesis and biochemistry assays. This effect aligns with the known mechanism of INO80, whose Ino80 motor engages nucleosomal DNA at the SHL-6 position near the H2A/mH2A C-terminal tail. In contrast, Chd1 binds exclusively at SHL2/-2 and is therefore unaffected by structural changes at the terminal DNA.

Stronger histone-DNA interactions around the entry/exit sites, such as those mediated by linker histone H1, are known to inhibit ATP-dependent chromatin remodeling by limiting access to flanking DNA required for remodeler engagement and nucleosome sliding (41). Our biochemical data suggest that the mH2A linker acts similarly, restricting linker DNA flexibility and accessibility, thereby hindering the ability of chromatin remodelers. The C-terminal tail of mH2A histone fold, on the other hand, does not alter entry/exit DNA stability but can inhibit remodeling by INO80 complex **(Fig. 6)**

Histone variants are known to also influence chromatin higher-order structures, although our understanding at this level remains limited compared to what is known at the mono-nucleosome level. In a previous cryo-EM study, we showed that reconstituted H2A.Z chromatin fibers adopt a more compact structure than canonical fibers *in vitro* (*26*). Interestingly, it was shown that histone variant H3.3 inhibit H2A.Z-mediated chromatin higher-order structure formation (42). A separate AUC study demonstrated that the mH2A linker promote self-association of nucleosome arrays *in vitro*, whereas the macro domain counteracts this compaction-promoting effect (14). How the mH2A linker and its macro domain interacts with linker DNA in compact chromatin domains, and how such interactions influence chromatin dynamics and higher-order folding, remains to be elucidated.

## DATA AVAILABILITY AND ACCESSION NUMBERS

The data that support the findings of this study are openly available in the Electron Microscopy Database (https://www.ebi.ac.uk/pdbe/emdb) and the Protein Data Bank (https://www.rcsb.org/). The EM map is deposited at the Electron Microscopy Database under accession code EMD-71726. The corresponding protein coordinate is deposited at the Protein Data Bank under PDB ID 9PM0.

## ACKNOWLEDGEMENT

We thank the scientists Dr. Liguo Wang, Dr. Guobin Hu, and Mr. Jake Kaminsky from the Laboratory for BioMolecular Structure (LBMS) for their help in data collection. LBMS is supported by the DOE Office of Biological and Environmental Research (KP1607011).

## AUTHOR CONTRIBUTIONS

V.S., R.J., A.M, G.L. prepared the samples for biochemistry and EM study. V.S., R.J., and A.M. performed the biochemical analysis and quantification with the assistance of G.L.; B.P. performed Alphafold3 structure prediction; D.T. performed image analysis and model building; D.T. oversaw the project and wrote the manuscript with help from all authors.

---

**Figure S1.**
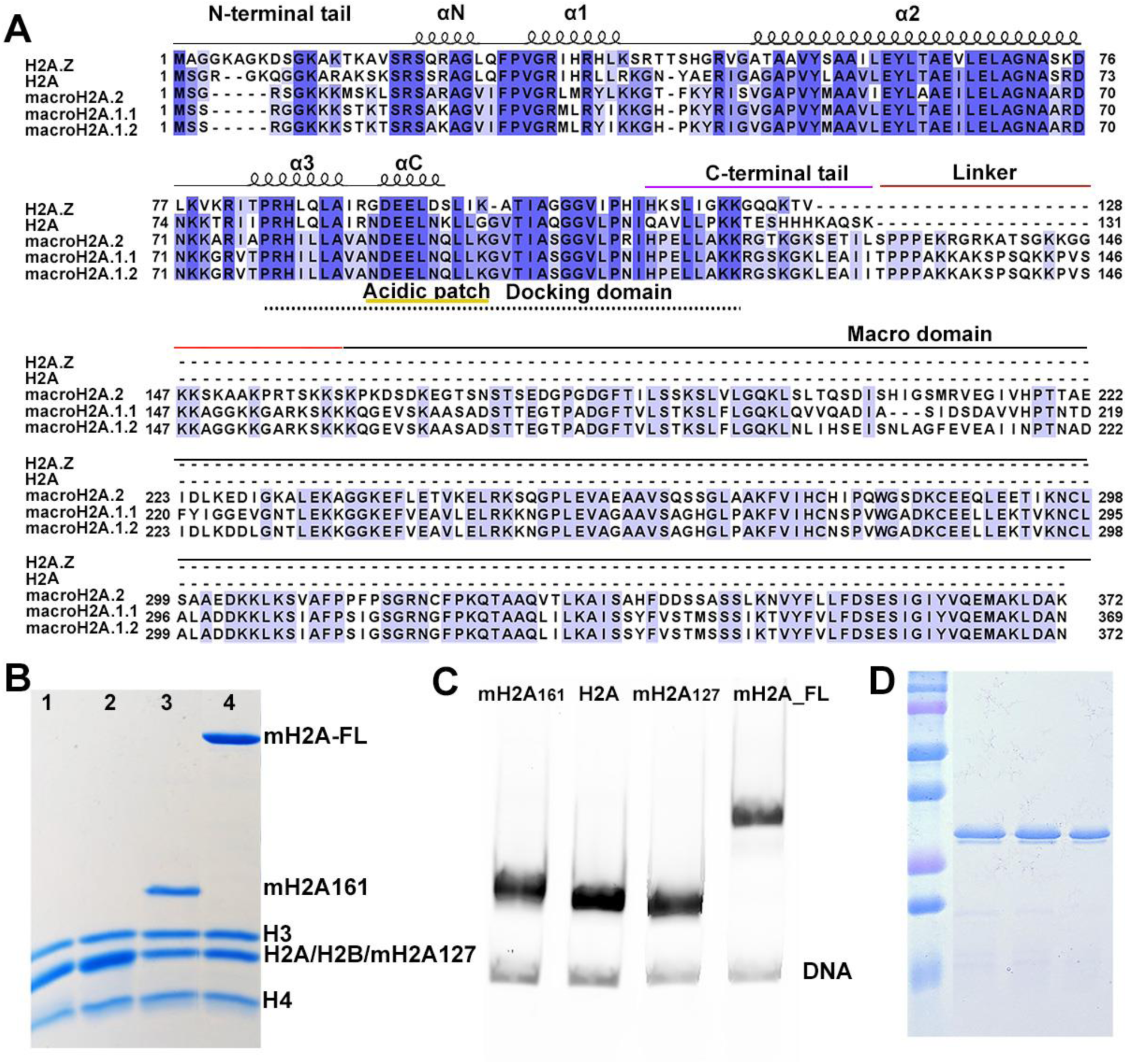
Sequence alignment and protein preparation. A. Sequence alignment of human H2A, H2A.Z, and mH2A. B. 15% of acrylamide SDS-PAGE gel show the histones in mH2A_127_ nucleosome (lane #1), H2A nucleosome (lane #2), mH2A_161_ nucleosome (lane #3), and mH2A-FL nucleosome (lane #4). C. 3% of native-PAGE gel shows the four different nucleosomes used in the study. D. 15% of acrylamide SDS-PAGE gel shows the recombinant scFv.

**Figure S2.**
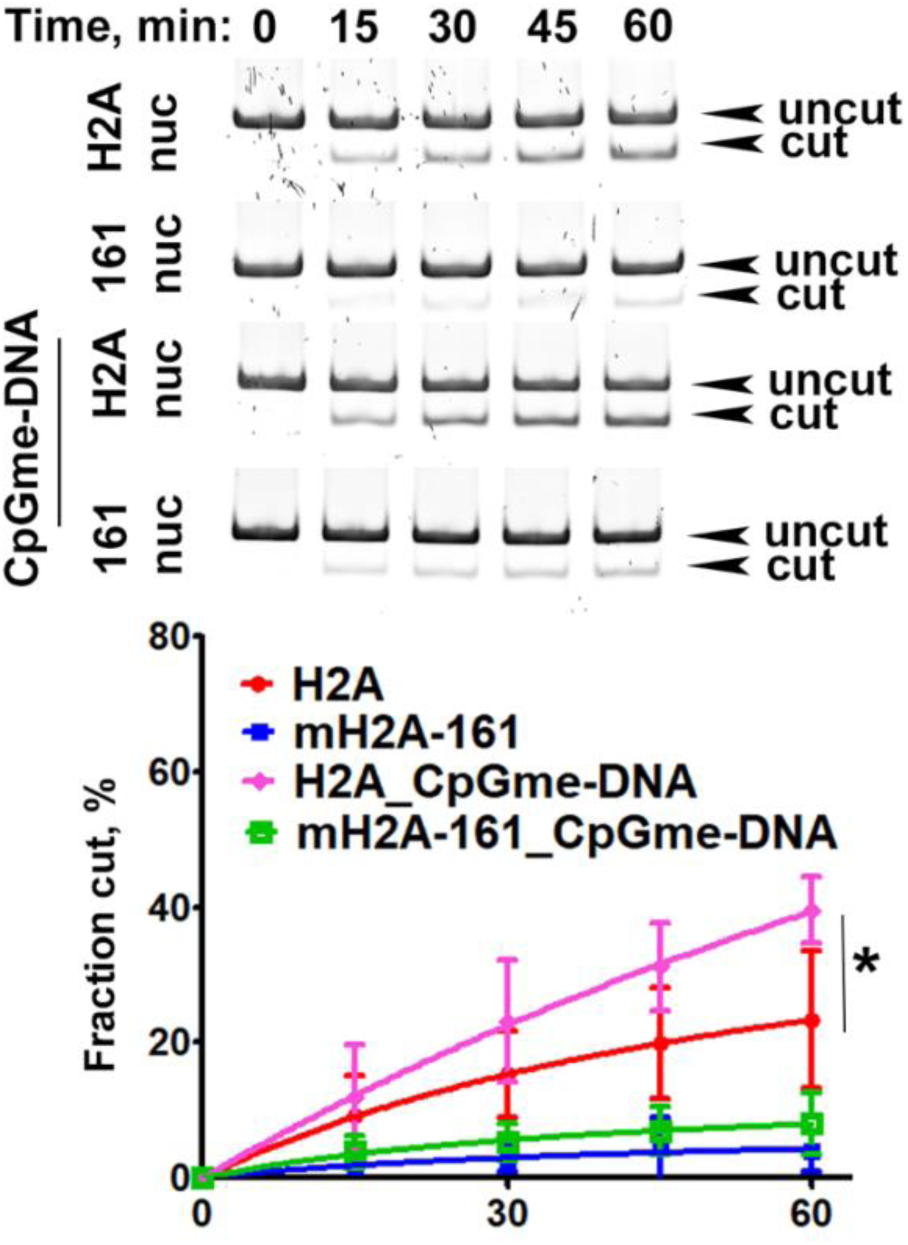
Restriction enzyme digestion of nucleosomes with methylated DNA. A representative acrylamide gels of HinfI digestion of canonical and mH2A_161_ nucleosomes reconstituted with and without CpG methylated DNA (top), and the quantification of the experiments. Data are mean ± SD, n = 3. Statistical significance was indicated by asterisk.

**Figure S3.**
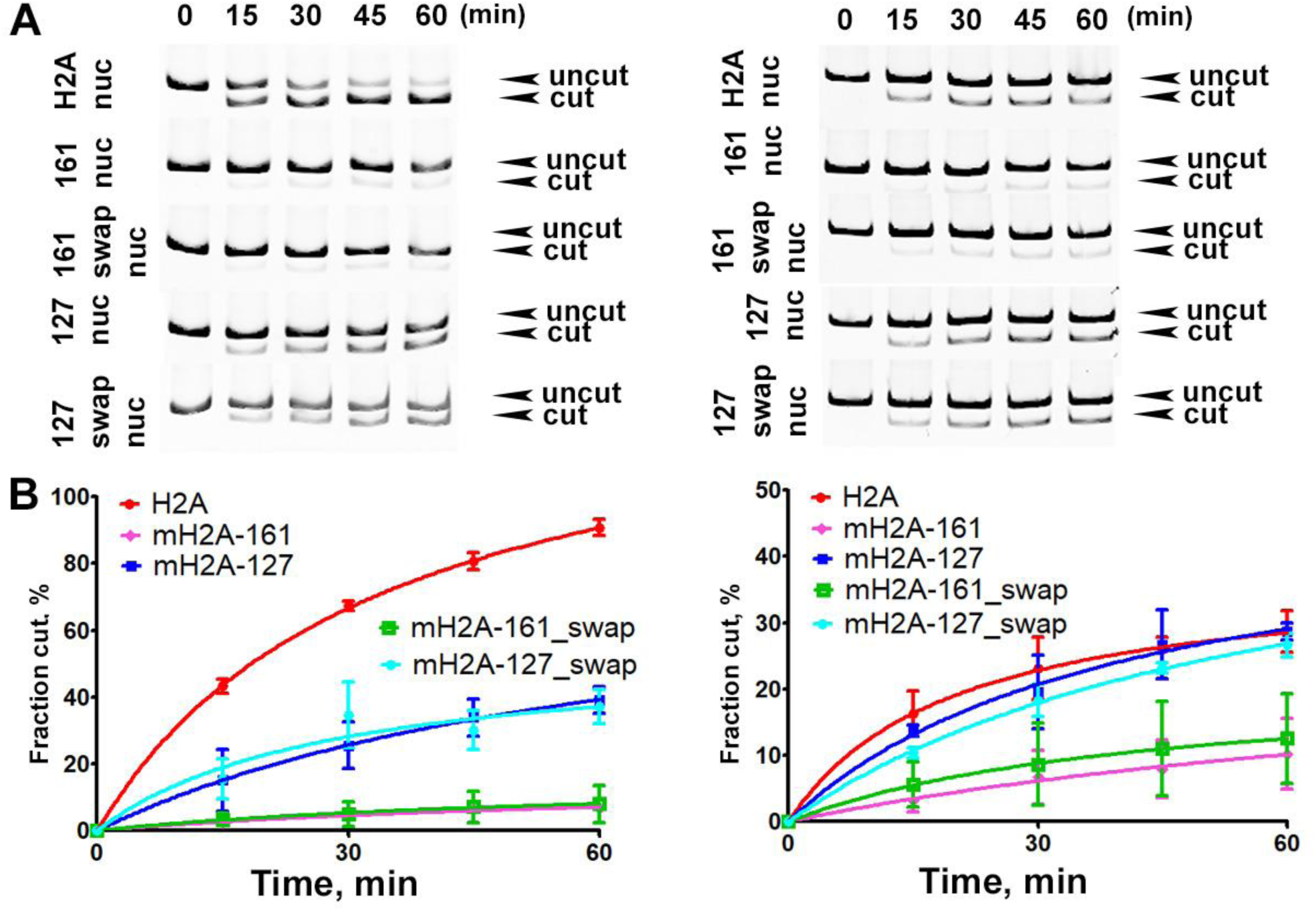
Restriction enzyme digestion of mH2A mutant nucleosomes. (A) Representative acrylamide gels show HinfI digestion (left) and BtsCI digestion (right) of H2A and various mH2A nucleosomes. (B) Quantification of the results in (A). The left panel is the HinfI digestion result and the right is BtsCI digestion results. Data are mean ± SD, n=3.

**Figure S4.**
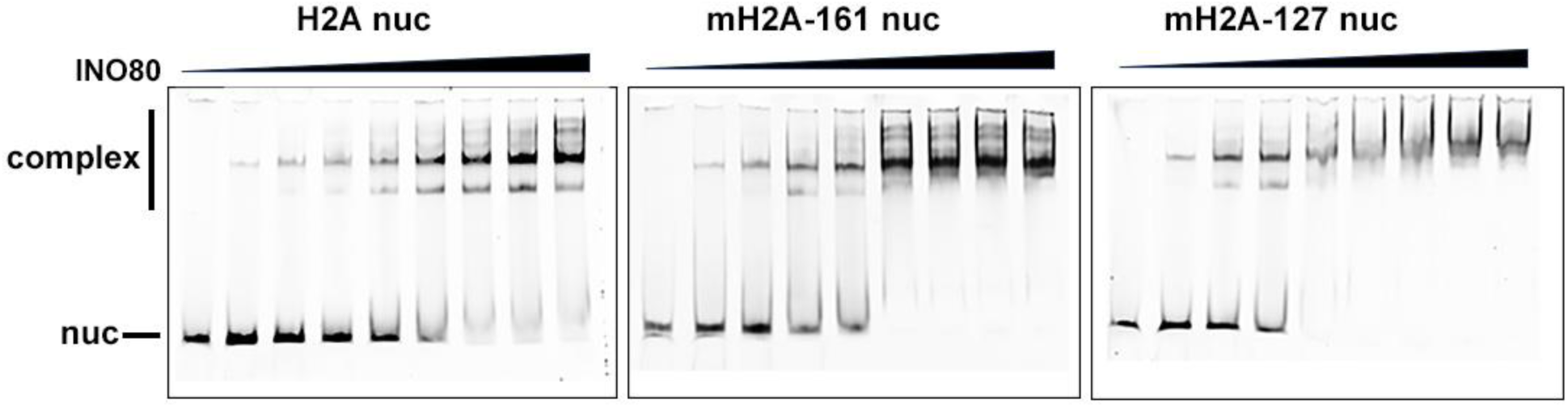
INO80 binds to H2A and various mH2A nucleosomes. 4% Native-PAGE shows complex formation of INO80 and nucleosomes. Each reaction contained 5 nM nucleosomes, while increased concentration of INO80 was used. The lanes in each gel (from left to right) correspond to 0, 50, 75, 100, 125, 150, 200, 225, and 250 nM INO80.

**Figure S5.**
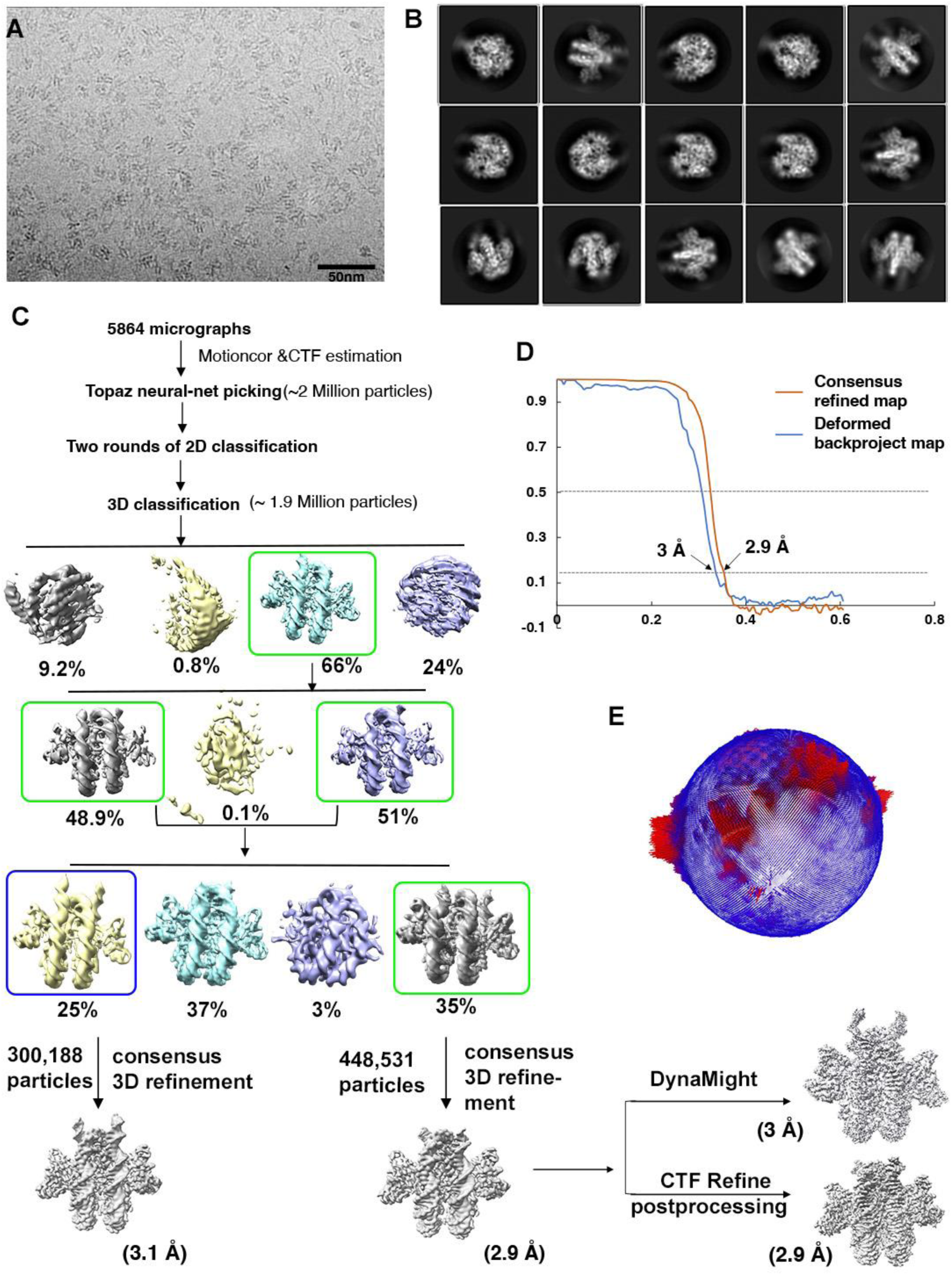
single-particle cryo-EM analysis of mH2A_161_ nucleosome with scFv. (A). Representative micrograph of vitrified sample. (B). Selected 2D class averages of mH2A_161_ nucleosomes. (C). single-particle cryo-EM workflow. (D). Fourier Shell Correlation (FSC) curve to show the estimated resolution of the density maps in this study. (E). Angular distribution of the consensus refined map.

**Figure S6.**
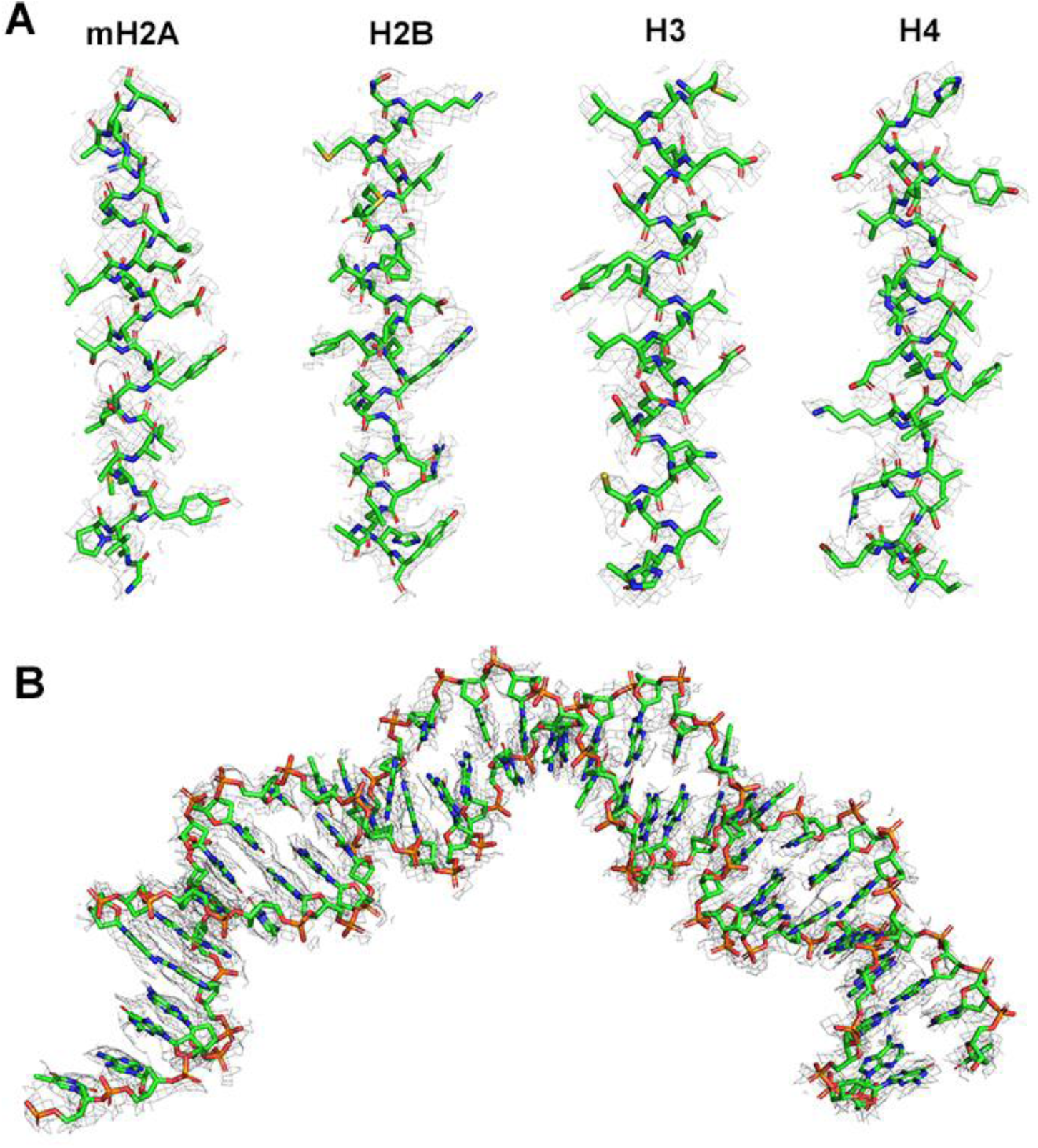
Fitting of molecular coordinates to selected secondary structure of the map to show the resolution. (A) The *α*2 helix of histone mH2A, H2B, H3, and H4. (B) A segment of DNA

**Figure S7.**
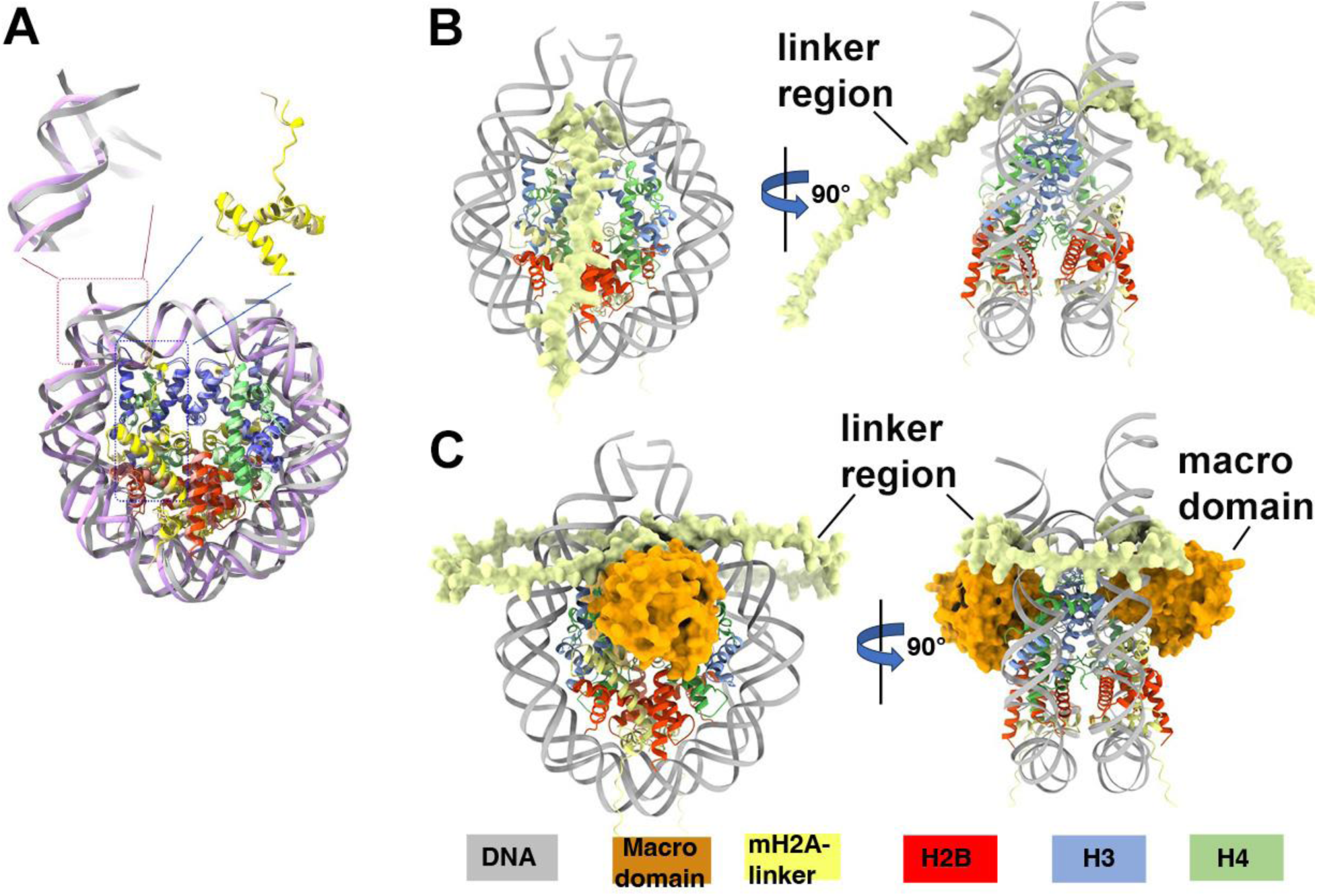
Structural comparison of mH2A nucleosome models. (A) Overlay of mH2A_161_ coordinate from this study with the crystal structure of mH2A nucleosome (PDB 1U35). One side of the entry/exit DNA and the C-terminus of the histone fold of mH2A are highlighted. (B) AlphaFold3 structure of mH2A_161_ nucleosome predicts that its linker regions (yellow) extend away from the nucleosome surface. (C) AlphaFold3 structure of mH2A-FL nucleosome shows the bended linkers (yellow) and the globular macro domains (orange).

