## Supplementary material for "MacroH2A-Mediated Gene Repression through Nucleosome Compaction and Remodeling Inhibition": sup-tables

**Supplemental Table 1 Cryo-EM data collection**

| Dataset | | mH2A_161_ nucleosome  (EMD-71726) |
| --- | --- | --- |
| **Data acquisition and processing** |  | |
| Microscope  Voltage (kV) | Krios  300kV | |
| Detector  Magnification | K3  105,000X | |
| Defocus range (µm)  Total dose (e/ Å^2^) | -0.9 to -2.5 µm  50 | |
| Symmetry | C1 | |
| Initial particle images (no.) | 1,281,469 | |
| Final particle images (no.) | 448,531 | |
| Map resolution (Å) | 2.9 | |
| FSC threshold | 0.143 | |
| Symmetry | C1 | |
| Map sharpening B-factor (Å) | -134 | |

**Supplemental Table 2 Model refinement and validation of mH2A_161_ nucleosome**

| Refinement | | mH2A_161_ nucleosome  (PDB ID 9PM0) |
| --- | --- | --- |
| Initial model used | Alphafold3 structure  PDB ID | |
| Model resolution (Å)  Model composition | 3.0 | |
| Protein residues  Nucleotides | 1224  300 | |
| R.m.s. deviations |  | |
| Bond length (Å) | 0.008 | |
| Bond angle (°) | 0.795 | |
| **Validation** |  | |
| Molprobity Score | 1.66 | |
| Molprobity clashscore | 4 | |
| Rotamer outlier (%) | 0.78 | |
| Cβ deviation (%) | 0.00 | |
| Ramachandran plot outlier (%) | 0.92 | |

**Supplemental Movie 1**. Consensus refined map of mH2A_161_ nucleosome

**Supplemental Movie 2**. DynaMight deformed reconstruction of mH2A_161_ nucleosome

**Supplemental Movie 3.** CryoSPARC 3D variability analysis volume series
